# Fine tuning cyclic-di-GMP signaling in *Pseudomonas aeruginosa* using the type 4 pili alignment complex

**DOI:** 10.1101/2020.10.17.343988

**Authors:** Shanice S. Webster, Calvin K. Lee, William C. Schmidt, Gerard C. L. Wong, George A. O’Toole

## Abstract

To initiate biofilm formation it is critical for bacteria to sense a surface and respond precisely. Type 4 pili (T4P) have been shown to be important in surface sensing, however, mechanism(s) driving downstream changes important for the switch to biofilm growth have not been clearly defined. Here, using macroscopic bulk assays and single cell tracking analyses of *Pseudomonas aeruginosa*, we uncover a new role of the T4P alignment complex protein, PilO, in modulating the activity of the diguanylate cyclase (DGC) SadC. Two hybrid and bimolecular fluorescence complementation assays show that PilO physically interacts with SadC and that the PilO-SadC interaction inhibits SadC’s activity resulting in decreased biofilm formation and increased motility. We show that disrupting the PilO-SadC interaction contributes to greater variation of cyclic-di-GMP levels among cells, thereby increasing cell-to-cell heterogeneity in the levels of this signal. Thus, this work shows that *P. aeruginosa* uses a component of the T4P scaffold to fine-tune the levels of this nucleotide signal during surface commitment. Finally, given our previous findings linking SadC to the flagellar machinery, we propose that this DGC acts as a bridge to integrate T4P and flagellar-derived input signals during initial surface engagement.

**Significance Statement:** T4P of *P. aeruginosa* are important for surface sensing and regulating intracellular cyclic-di-GMP levels. This work identifies a new role for the T4P alignment complex, previously known for its role in supporting pili biogenesis, in surface-dependent signaling. Furthermore, our findings indicate that *P. aeruginosa* uses a single DGC, via a complex web of protein-protein interactions, to integrate signaling through the T4P and the flagellar motor to fine-tune cyclic-di-GMP levels. A key implication of this work is that more than just regulating signal levels, cells must modulate the dynamic range of cyclic-di-GMP to precisely control the transition to a biofilm lifestyle.

## Introduction

Biofilms are surface-attached multi-cellular communities and a key early step in biofilm formation is surface sensing [1]. Flagella and type 4 pili (T4P) are required for detecting surface contact and for surface signaling [2–5]. For example, the T4P are thought to act as a “force transducer” by detecting the resistance to retraction when cells are surface engaged, thereby activating downstream pathways such as synthesis of holdfast in *Caulobacter* or activation of the Chp chemosensory [2, 3] and FimS-AlgR two component system [6] in *P. aeruginosa*.

Despite multiple studies implicating the T4P [2, 3, 7] and flagella [8, 9] in surface sensing, the mechanism(s) whereby these motility machines transduce and integrate such signals has not been established. Here we show that *P. aeruginosa* uses a T4P alignment complex protein, PilO, to interact with the diguanylate cyclase (DGC) SadC to sequester and reduce the activity of SadC, which results in decreased surface motility and reduced biofilm formation. We show that disrupting the PilO-SadC interaction results in greater variation in cyclic-di-GMP levels among signaling cells; thus, this complex regulatory network seems necessary to maintain a uniform output of this dinucleotide signal in a given signaling population. Finally, given the documented role of SadC in interacting with a component of the flagellar motor [10], we propose a model whereby this DGC can act as a bridge to integrate surface-derived input signals from both the T4P and flagella.

## Results

### Type 4 pili alignment complex protein, PilO, physically interacts with SadC

We previously reported genetic studies supporting the model that the T4P PilMNOP proteins (i.e. the pilus alignment complex) are involved in signal transduction from the cell surface protein, PilY1, to inner membrane-localized DGC SadC [6] (**Fig. 1A**), but the mechanism underlying this signaling has not been defined. By leveraging the observation that PilY1 protein levels increase when cells are surface-grown, we showed that repression of swarming motility did not occur when PilY1 was overexpressed in the *ΔpilMNOP* mutant or for strains carrying single non-polar mutations in the *pilM, pilN, pilO* or *pilP* genes suggesting a role for the PilMNOP proteins in signaling from PilY1 [6]. Similarly, a strain carrying a *sadC* deletion also lost PilY1-dependent swarm suppression [11], implicating the alignment complex and SadC in PilY1-mediated, surface-dependent control of cyclic-di-GMP.

**Figure 1.**
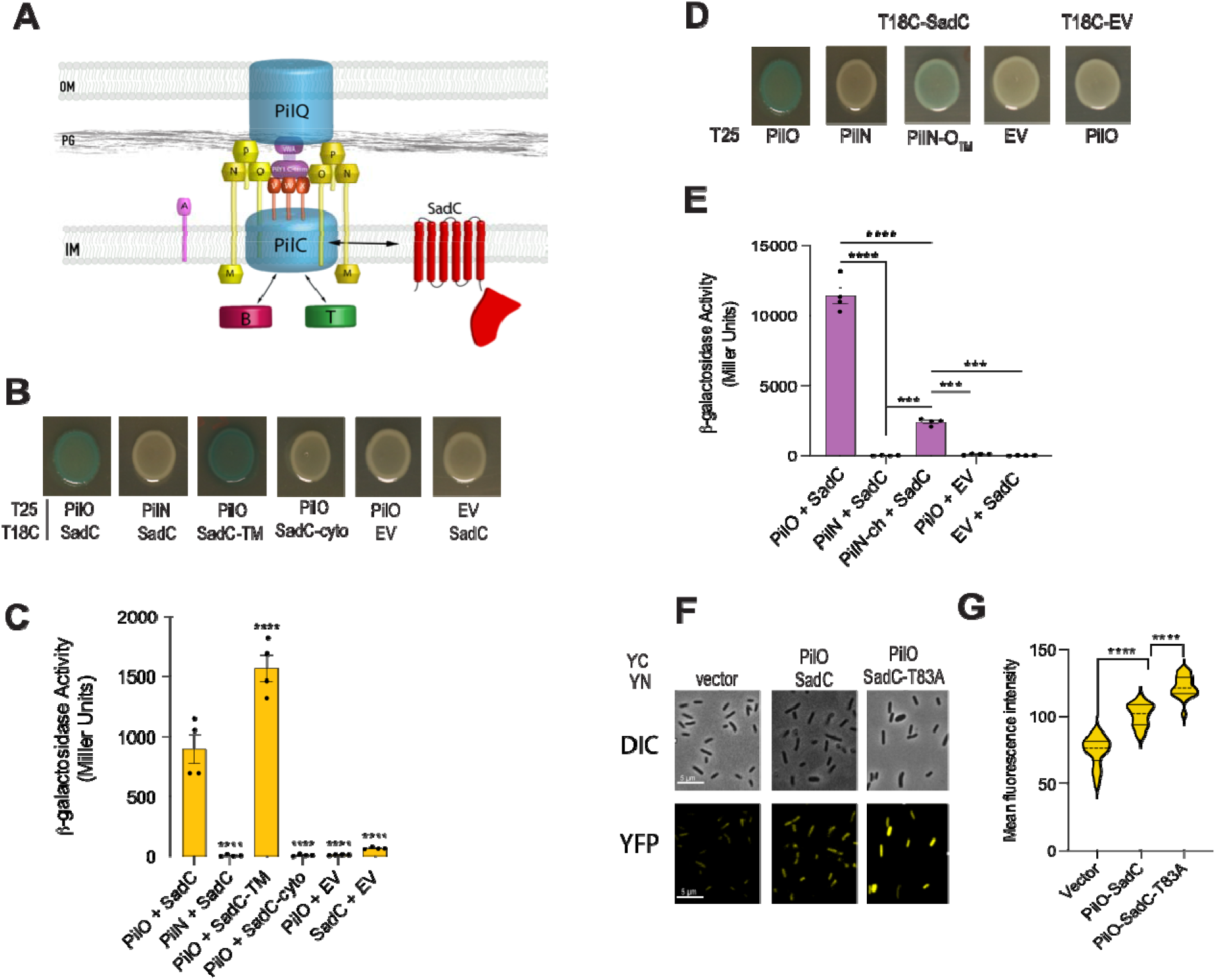
Physical interaction between PilO and SadC is mediated by their TMD. **A.** Shown is a model of the T4P machinery, featuring SadC interaction with PilO, a component of the T4P alignment complex. Legend: outer membrane (OM), inner membrane (IM), and peptidoglycan (PG). T4P apparatus is highlighted in blue: the baseplate PilC, the secretin PilQ, the major pilin PilA, and the minor pilins PilVWX. The alignment complex (PilMNOP) is labeled in yellow. The extension and retraction ATPases, PilB (red) and PilT (green) are also shown. **B.** PilO interacts with SadC by BACTH assay. Images of spots of co-transformations with the indicated proteins fused to the C-terminus of the T25 or T18 domains of adenylate cyclase following incubation at 30°C for 40 h and then at 4°C for three days to allow for further color development on X-gal-containing plates. Empty vectors are the negative controls in this and subsequent experiments. **C.** Beta-galactosidase activity in Miller units for interactions shown in **B. D.** Images from BACTH analysis for SadC co-transformed with PilO, PilN or PilN-O™ chimera. PilN-O™ is a chimeric protein of PilN with its transmembrane domain replaced with that of PilO. **E.** Quantification of beta-galactosidase activity of co-transformation from **D** shown in Miller units. Beta-galactosidase activity in panels **C** and **E** were quantified from cells scraped from transformation plates supplemented with antibiotics and X-gal. Error bars show SEM from four biological replicates and statistical significance was determined using one-way analysis of variance (ANOVA) and Dunnets multiple comparison post-hoc test. p-values: p ≤ *** 0.0001, p ≤ **** 0.0001. **F**. PilO-SadC interaction shown by bimolecular fluorescence complementation (BiFC) analysis. DIC (top) and fluorescent (bottom) images from BiFC assay shown for the vector only control, and PilO with either WT SadC or the SadC-T83A variant. The N-terminus of PilO and SadC proteins were fused to the C-terminal (YC) and N-terminal (YN) portions of the yellow fluorescent protein (YFP), respectively. Representative images of fluorescence intensity for interaction pairs as indicated. The vector is included as the negative control. **G**. Quantification of mean fluorescence intensity per cell. Dashed lines on violin plots represent the median and solid lines represent the first and third quartiles. Data points are the mean fluorescence intensity per cell from at least six fields. Data are from two independent experiments on different days, with the P-value calculated from a Mann-Whitney U test. p-values: p ≤ ***0.001, p ≤ ****0.0001

The PilMNOP alignment complex is stabilized by a series of documented protein-protein interactions between PilP-PilN, PilP-PilO, PilN-PilM and PilN-PilO that spans the cytoplasm, across the inner-membrane to the periplasm of the cell [12, 13]. Based on our prior findings and the known interactions among the PilMNOP proteins, we hypothesized that physical interaction between SadC and one or more components of the T4P alignment complex might be important in modulating SadC-dependent cellular cyclic-di-GMP levels, which would in turn affect biofilm formation and motility. We focused on PilN and PilO because these proteins share the inner membrane-localization of SadC [12, 14]. Using bacterial adenylate cyclase two hybrid (BACTH) we show a significant interaction between PilO and SadC but not with the structurally similar PilN (**Fig. 1B and C**), suggesting that the interaction between PilO and SadC is specific.

We next sought to define which portions of SadC and PilO interact. SadC has 6 predicted membrane helices at its N-terminus constituting a transmembrane domain (TMD) and a C-terminal cytoplasmic GGDEF catalytic domain, while PilO has a single TMD, and an extended periplasmic domain (**Fig. 1A**). We tested interactions between the transmembrane or the GGDEF domain of SadC with full length PilO in the BACTH system. PilO interacts with the TMD of SadC, while there was a lack of interaction with the SadC’s catalytic domain (**Fig. 1B** and **1C**).

Given that PilO has a periplasmic globular head domain and a 22 amino acid α-helix that extends into the inner membrane (IM), we hypothesized that the TMD of PilO is important for interaction with the transmembrane of SadC. To evaluate this prediction, we constructed a chimeric protein with the periplasmic domain of PilN and the TMD of PilO (amino acids 28-49), which we designated PilN-PilO™. BACTH analysis showed significantly more interaction between the PilN-PilO™ chimera and SadC compared to full length PilN and SadC (**Fig. 1D** and **1E**). However, we do not observe as much interaction with the chimeric protein as we observed for full length PilO suggesting that other parts of PilO may be important for interacting with SadC.

As a second method to assess the PilO-SadC interaction, we used bimolecular fluorescence complementation (BiFC). Cells expressing both PilO and SadC fused to the N-terminal and C-terminal halves of the yellow fluorescent protein (YFP) showed a robust fluorescent signal as compared to the vector control (**Fig. 1F** and **1G**).

### Surface-exposed residues of SadC TMD modulate interaction with PilO

Given that PilO is able to interact with SadC, we hypothesized that mutations in the TMD of SadC could modulate interaction with PilO. To test this hypothesis, we performed BACTH analysis coupled with a genetic screen using error-prone PCR mutagenesis of the TMD of SadC and screened for mutants that affected interaction with PilO. From this screen, we identified two candidate mutations, SadC-T83A and SadC-L172Q. We re-tested candidates in the BACTH system and found that the SadC-T83A variant increases interaction while the SadC-L172Q mutant protein disrupts interaction with PilO (**Fig. 2A** and **2B**). BiFC analysis of SadC-T83A mutant allele showed increased fluorescence as compared to the vector control (**Fig. 1F** and **1G**), corroborating the results from the BACTH.

**Figure 2.**
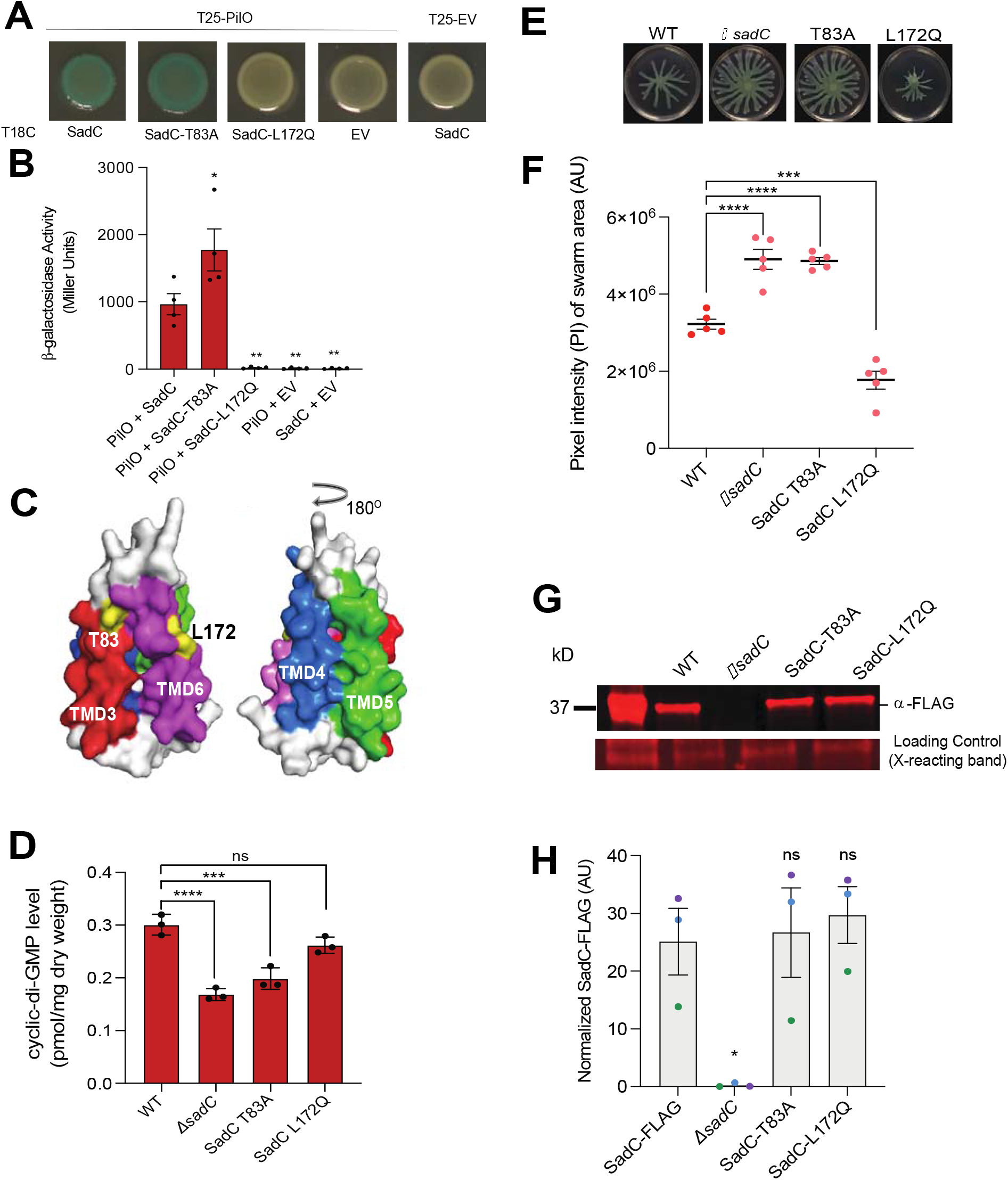
Mutations in SadC’s TMD modulate interaction with PilO and impact cyclic-di-GMP-associated behaviors in *P. aeruginosa*. **A.** Images from BACTH analysis for co-transformations with PilO and either WT SadC, SadC-T83A and SadC-L172Q proteins. **B.** Quantification of beta-galactosidase activity in Miller units for interactions in **A**. Details of experiments provided in the legend of Figure 1. **C.** Predicted structure of four of the six N-terminal TMD (amino acids 1-187) of SadC. The structure was generated using the prediction software Phyre [20]. TMD3 (red), TMD4 (blue), TMD5 (green) and TMD6 (magenta) are shown. Residues T83 and L172 located on TMD3 and TMD6, respectively, are highlighted in yellow and labeled. **D**. Quantification of global cyclic-di-GMP levels for WT and *sadC* variants. **E.** Representative swarm images. **F.** Quantification of pixel intensity (PI) of swarm area for images shown in **E**. Error bars in **B, D** and **F** are SEM and statistical significance was determined by one-way ANOVA and a Dunnets post-hoc test, p-values: p ≤ * 0.01, p ≤ ** 0.001, p ≤ *** 0.0001 and p ≤ **** 0.00001; ns, not significant. **G.** Representative blot for normalized SadC-3xFLAG protein levels. The band at ~30 kD is a non-specific, cross-reacting band with the anti-FLAG antibody and serves as an additional loading control. **H.** Quantification of normalized SadC-FLAG protein levels in WT and mutants relative to the cross-reacting band. Data are from three biological replicates. Error bars are SEM and statistical significance was determined by one-way ANOVA and a Dunnets post-hoc test. p-values: p ≤ * 0.05, ns, not significant.

Using the prediction server Phyre [15], we generated a model of the SadC TMD with an 80% confidence interval based on homology to KdpD, a sensor protein with 6 membrane helices and a member of the two-component KdpD/KdpE regulatory system of *Escherichia coli*. The TMD regions of the KdpD and SadC share 40% sequence identity. The T83 and L172 residues of SadC were mapped to TMD3 and TMD6, respectively. Despite their mapping to distinct TMD, these mutations map to the surface and the same face of this model (**Fig. 2C**), suggesting that these surface-exposed, proximal residues of SadC both participate in the interaction with PilO.

### Mutations in the SadC TMD that impact interaction with PilO have functional consequences in *P. aeruginosa*

To determine if the mutations in SadC that modulate interaction with PilO impact *P. aeruginosa* surface behaviors we introduced the point mutations onto the chromosome. The SadC-T83A mutant hyper-swarmed and showed decreased biofilm levels as compared to WT (**Fig. 2D-F** and **Fig. S1**), indicating reduced c-di-GMP levels. The level of cyclic-di-GMP in the strain expressing the SadC-T83A mutant protein is not significantly different from the *ΔsadC* strain (**Fig. 2D**). Consistent with this observation, the strain expressing the SadC-T83A mutant protein and the *ΔsadC* deletion showed the same hyper-swarming and reduced biofilm phenotypes (**Fig. 2D-F** and **Fig. S1**). In contrast, the strain carrying the SadC-L172Q mutant protein showed reduced swarming motility (**Fig. 2E-F**), but a non-significant change in global cyclic-di-GMP levels (**Fig. 2D**) and no significant change in early biofilm formation (**Fig. S1**). These data indicate that disrupting the PilO-SadC interaction is not sufficient to activate SadC, a point we discuss further below.

To ensure that the observed phenotypes caused by these mutations in SadC were not due to differences in steady state protein expression levels, we performed Western blot analysis on FLAG-tagged SadC mutant variants and showed that SadC-T83A and SadC-L172Q variants were as stable as WT FLAG-tagged SadC (**Fig. 2G** and **2H**).

These data, together with the BACTH and BiFC analyses, indicate that mutations that increase interaction between PilO and SadC decrease cellular levels of cyclic-di-GMP and suggests a model wherein PilO sequesters SadC and inhibits its activity.

### The Small-xxx-Small motif in the PilO transmembrane mediates interaction with SadC

Given that PilO interacts with SadC via its TMD, we wanted to determine whether there was a specific motif present in the transmembrane of PilO that might be important for interaction with SadC. We scanned the PilO transmembrane and found that there is a conserved Small-xxx-Small motif present at residues 40 and 44 (A40xxxA44). This motif has been shown to play an important role in helix-helix dimerization in transmembrane proteins [16, 17], thus we hypothesized that this domain might be mediating interaction between the α-helix of PilO and the SadC TMD. We mutated the A40 and A44 residues to glutamate (E40xxxE44) and tested this mutant protein for interaction with SadC using BACTH assay. PilO-ExxxE mutation disrupts interaction with SadC (**Fig. 3A**). However, the PilO-ExxxE mutant rendered the protein less stable and decreased IM protein levels of PilO as compared to WT PilO (**Fig. 3B-D**), therefore it is possible that the decreased interaction observed is due, at least in part, to reduced IM-localized PilO-ExxxE protein.

**Figure 3.**
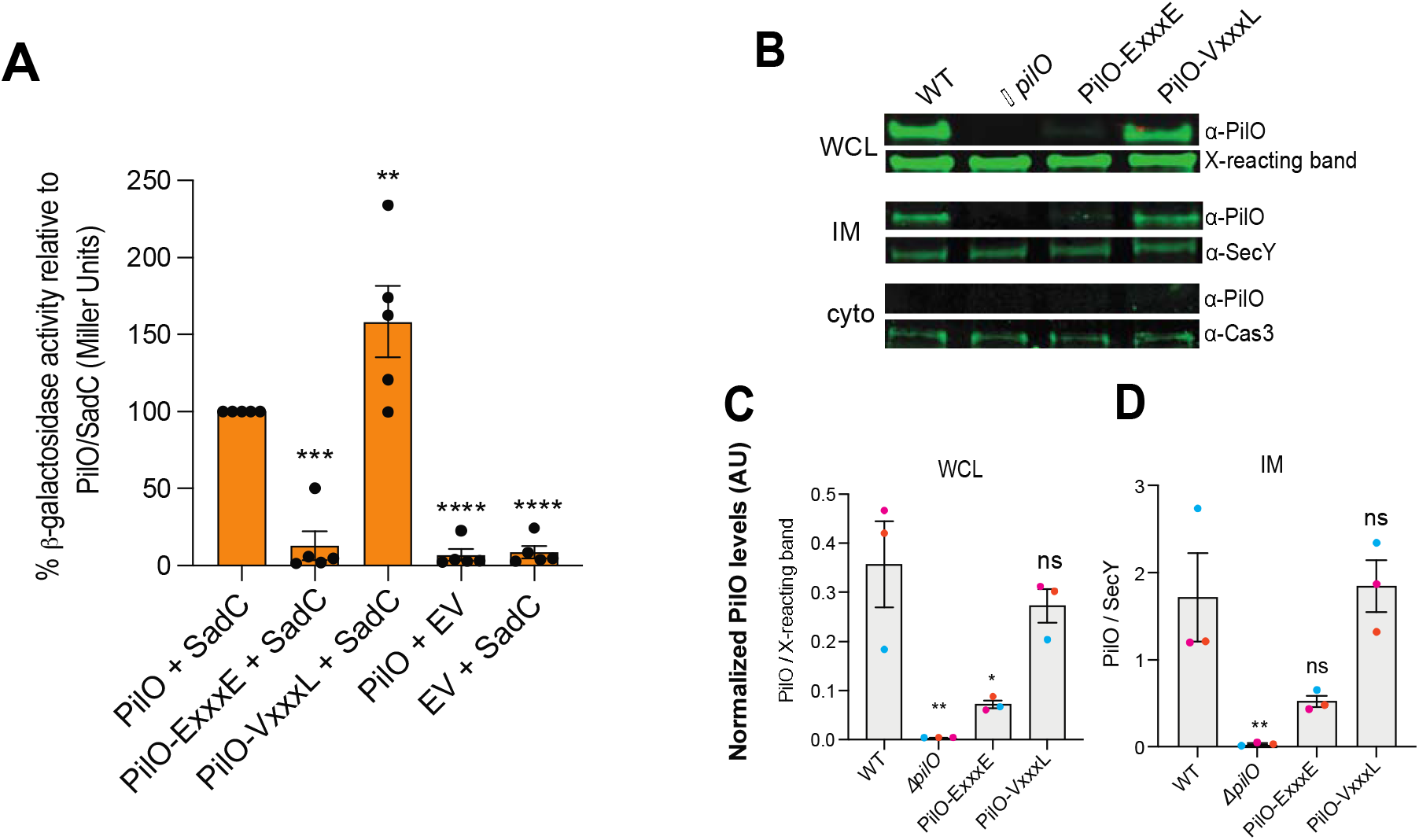
A conserved Sm-xxx-Sm motif in the TMD of PilO is important for interaction with SadC. **A.** Beta-galactosidase activity from BACTH assay for SadC co-transformed with PilO, PilO-ExxxE and PilO-VxxxL mutant proteins. The percent beta-galactosidase activity relative to PilO-SadC (set to 100%) is shown. Dots shown on graph for beta-galactosidase assay represent data points from five biological replicates. Error bars are SEM, and statistical significance was determined by a one-way ANOVA and a Dunnets post-hoc test. p-values: p ≤ ** 0.001, p ≤ *** 0.0001, p ≤ **** 0.0001. **B**. Representative images for PilO protein levels in whole cell lysate (WCL), inner-membrane (IM) and cytoplasmic (cyto) fractions for WT and PilO variants. PilO was not detected in the cytoplasmic fraction. The cytoplasmic protein, Cas3 (~120 kD) was used as a loading control for the cytoplasmic fraction. **C** and **D**. Quantification of normalized PilO protein levels in WCL and IM, respectively. PilO protein levels in WCL were normalized to a cross-reacting band (X-reacting band) at ~43 kDa while IM levels were normalized to the IM localized protein, SecY, ~50 kDa (**C**). Data in panels **C** and **D** represent three biological replicates. p-values from a one-way ANOVA with a Dunnet post-test, p-values: p ≤ * 0.05, p ≤ ** 0.01; ns, not significant.

As an alternative approach to identify mutants in the A40xxxA44 motif of PilO that impact PilO-SadC interaction, we performed a targeted screen wherein we used a primer-based approach to introduce random mutations at amino acid positions 40 and 44 in the PilO-containing BACTH fragment to generate a mutant library. We then tested the PilO mutant library for interaction with SadC in the BACTH system and screened for variants of PilO that increased interaction with SadC, as judged by dark blue colonies in the BACTH assay. We sought mutants that enhanced interaction because we postulated that these alleles were likely to be stable. From this screen, we identified a candidate mutant, PilO-VxxxL. BACTH studies confirmed that the PilO-VxxxL mutant protein interacts significantly more strongly with SadC than does the WT PilO (**Fig. 3A**). PilO-VxxxL shows similar levels of protein expression and IM localization as WT in whole cell lysates and IM fractions, respectively (**Fig. 3B-D**). Furthermore, the strain expressing the PilO-VxxxL mutant protein twitches to the same extent as WT, which demonstrates that these mutations do not disrupt the key role of this protein in the T4P alignment complex (**Fig. S2A**).

Given the findings for the SadC-T83A protein variant that enhance interaction with PilO and results in decrease biofilm formation, hyper-swarming and reduce global cyclic-di-GMP levels, we hypothesize that the PilO-VxxxL variant would cause similar phenotypic outputs. Surprisingly, however, using bulk assays we did not observe any significant changes in intracellular levels of cyclic-di-GMP, swarming motility, or biofilm formation as compared to WT for the PilO-VxxxL variant protein (**Fig. S2B-D**).

SadC interacts with both PilO and MotC, and the interaction between MotC and SadC stimulates SadC’s activity [10]. Thus, one simple explanation for the lack of bulk phenotypes for the strain carrying the PilO-VxxxL variant is that despite the increased interaction between SadC and PilO-VxxxL (which should reduce cyclic-di-GMP level based on our model), SadC’s ability to interact with MotC could mitigate the increased PilO-SadC-VxxxL interaction, thus causing phenotypes to be more subtle. We confirmed the SadC-MotC interaction, and we observed that SadC interacts more strongly with MotC than it does with PilO (**Figure S3A**), consistent with the idea that PilO and MotC may be competing for SadC binding. Furthermore, we demonstrated that the SadC-T83A and SadC-L172Q variants still interact with MotC at WT levels (**Figure S3B**), indicating that the PilO-SadC and the SadC-MotC interaction faces are distinct. We address the issue of phenotypes for the PilO-VxxxL mutant directly using single cell tracking below.

### PilO mutations that modulate interaction with SadC affect variation of cyclic-di-GMP signaling during surface growth

Our analysis above is consistent with PilO and SadC interacting, and this interaction between the proteins modulating SadC’s DGC activity. As mentioned above, however, some of the macroscopic bulk phenotypes of the strains carrying PilO mutations were unexpected, being either quite subtle or not significantly different from WT. To address this issue directly, we used single-cell tracking to investigate changes in cyclic-di-GMP levels during surface colonization by the WT and strains carrying mutant variants of the PilO and SadC proteins to explore how these mutations might impact cyclic-di-GMP production for populations of cells resolved at the single cell level.

We tracked single cell levels of cyclic-di-GMP in the WT and mutant strain using a reporter wherein GFP was fused to the cyclic-di-GMP-responsive P_cdrA_ promoter. We tracked single cells in a flow cell immediately before and after the initiation of exponential surface cell growth, as reported [18–20], and aggregated the data across 5 blocks of time of ~6 hrs each (**Fig. S4**). We noticed two populations of cells – those expressing and those not expressing GFP, thus a GFP signal cut-off was defined to partition cells into two these sub-populations of cyclic-di-GMP “on” and “off” states [21] (**Fig. S5**). The fraction of cyclic-di-GMP “on” cells in a population is a good measure for whether SadC is active or inactive in those cells, since a cyclic-di-GMP “on” cell should have active SadC producing c-di-GMP, and vice versa. For the WT and the PilO-VxxxL allele, which enhances PilO-SadC interaction, we observe a reduction in, and the leveling off of the number of cells of the cyclic-di-GMP “on” sub-population over time (**Fig. 4A** and **4B**). In contrast, the PilO-ExxxE, which disrupts the PilO-SadC interaction, or for a strain wherein PilO is deleted (Δ*pilO*), we observe the opposite pattern – a progressive increase in the cyclic-di-GMP “on” sub-population over time. These data are consistent with the model that PilO-SadC interaction is involved in controlling cyclic-di-GMP levels by affecting whether SadC is active or inactive. WT and increased PilO-SadC interactions lead to similar levels of cells with active SadC, as indicated by the similar trend of the fraction of cyclic-di-GMP “on” cells. However, disruption of this SadC-PilO interaction leads to an increasing fraction of cyclic-di-GMP “on” cells and thus more cells with active SadC.

**Figure 4.**
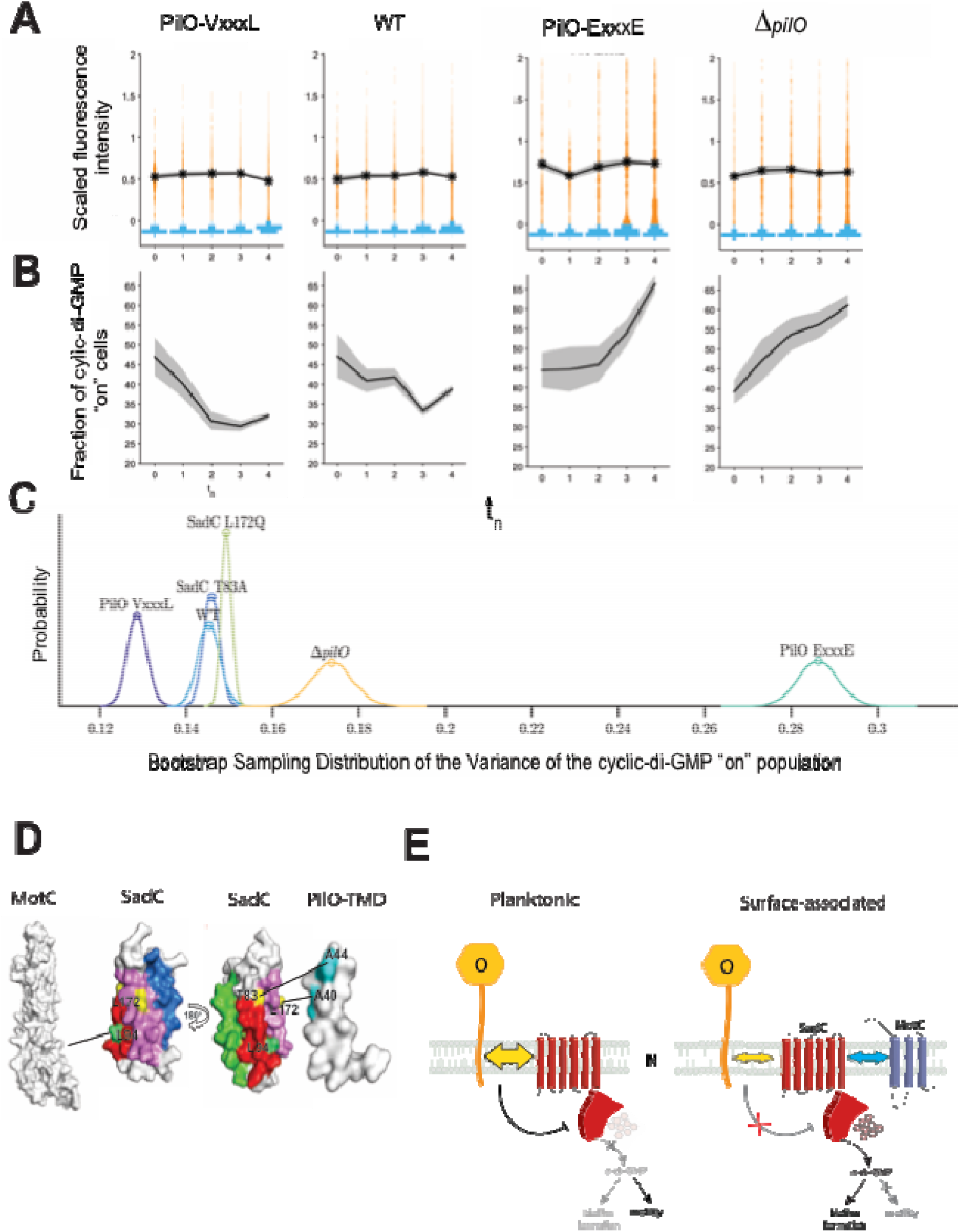
PilO-SadC interaction mutants analyzed by single cell tracking. **A.** GFP intensity was determined on a cell-by-cell basis for strains carrying the *P_cdrA_-gfp* construct, a reporter of cyclic-di-GMP levels. The GFP on/off cutoff is defined as outlined in Supplemental Figure S5. The yellow dots show the individual data points of the “on” population. The asterisks and lines show the mean intensity of the “on” population and the shaded area represents the 95% confidence interval, which is a reflection of the variation of the data used to generate the mean. The blue indicates intensity data from the “off” populations. The times indicated on the x axis are ~6 h time periods starting just prior to and at the onset of exponential attachment of bacteria to the flow cell (~20 hrs; see Suppl. Fig. 4). **B.** Shown is the fraction of the cyclic-di-GMP “on” population for each of the time periods in (A). The dark line is the fraction of “on” cells and the gray represents the 95% confidence interval. The times indicated on the x axis are ~6 h time periods starting just prior to and at the onset of exponential attachment of bacteria to the flow cell (~20 hrs; see Suppl. Fig. 4). **C**. Bootstrap sampling distributions of the variance of the entire cyclic-di-GMP “on” population (summing up all time points in panel A). Distribution overlap determines the p-value indicating whether the variance is significantly different. All strains have a significantly different variance compared to WT (p < 1e-4), except for SadC T83A (p = 0.43) and SadC L172Q (p = 0.08). The number of cells used to generate the distribution for each time point and strain is summarized in Table S1 of the Supplemental Material. **D.** Surface representation of the structures for MotC, SadC and PilO TMD determined using Phyre. TMDs of SadC colored as in Figure 2. SadC-L94P that disrupt interaction with MotC is shown in green while SadC mutations (T83A and L172Q) that modulate interaction with PilO are shown in yellow. The PilO TMD with the alanine residues of the Sm-xxx-Sm motif are shown in magenta. Rotation of SadC (180°) is shown to better view TMD3 with T83A and L94P residues. Location of SadC-L94P and SadC-T83A on the same alpha helix suggest that SadC does not simultaneously interact with both PilO and MotC. **E.** Proposed model for the role of the PilO-SadC interaction during transition from planktonic to surface-associated or a biofilm state. In a planktonic state, PilO-SadC interaction inhibits SadC’s activity which results in decreased biofilm formation and increased motility. During surface association PilO-SadC interaction is disrupted relieving repression of SadC activity; SadC in turn is activated through interaction with MotC, resulting in increased cyclic-di-GMP levels, which promotes biofilm formation and inhibits motility.

We see that PilO-SadC interactions can turn SadC “on” and “off.” However, there are additional surprising consequences of altering PilO-SadC interactions. Here, we focus only on the cyclic-di-GMP “on” subpopulation of cells (and thus cells with active SadC) and look at their GFP intensities, which are related to the level of cyclic-di-GMP production and thus the level of SadC activity. Interestingly, the cyclic-di-GMP “on” subpopulation had a broad distribution of intensities rather than a narrow range of intensities for all cells. This distribution of intensities reflects the heterogeneous levels of SadC activity across cells in a population. We can quantify this heterogeneity in SadC activity by calculating the variance of the cyclic-di-GMP “on” subpopulation of GFP intensities (**Fig. 4C**). A population with higher variance will have a wider range of cells with both high and low cyclic-di-GMP production and SadC activity, while a population with lower variance will have cells with a more uniform cyclic-di-GMP production and SadC activity. Interestingly, there exists an inverse trend between the variance of the cyclic-di-GMP “on” subpopulation (and SadC activity) and the strength of PilO-SadC interactions. Mutations that decrease the PilO-SadC interaction show higher variances in cyclic-di-GMP levels compared to WT, while mutations that increase the PilO-SadC interaction show similar or lower variances in cyclic-di-GMP levels compared to WT (**Fig. 4C**). Thus, these data suggest that PilO-SadC interactions also serve to enforce uniformity of SadC cyclic-di-GMP output by maintaining dinucleotide levels within a narrow window. Thus, increasing this PilO-SadC interaction leads to SadC activity being more uniform, while decreasing this interaction leads to SadC activity being more heterogeneous.

Taken together, the data suggest that PilO-SadC interactions are involved in maintaining cyclic-di-GMP homeostasis via two distinct but complementary modes: by regulating whether cyclic-di-GMP is produced or not (by controlling the active state of SadC), and by regulating the level of cyclic-di-GMP production (by tuning the activity of SadC). Weakening this interaction results in an impaired capacity to control the cyclic-di-GMP levels in a given population. The observed heterogeneity in cyclic-di-GMP production and SadC activity in a given population comes from both modes operating simultaneously as the cells maintain cyclic-di-GMP homeostasis.

## Discussion

Here we show that the PilO-SadC interaction is important for driving early steps in biofilm formation. How does this study fit in terms of an overall model for the transition from planktonic to biofilm state? When in a planktonic state, cells can freely extend and retract their T4P. In this state we propose that in a planktonic state PilO interacts with SadC to sequester SadC and inhibits the activity of this DGC. In contrast, once cells adhere to a surface, we propose that PilO’s interaction with SadC is reduced, and SadC then becomes activated, likely via its interaction with MotC [10], to increase cyclic-di-GMP levels (**Fig. 4D** and **4E**).

There are several important implications of our findings. First, our work shows that the T4P alignment complex has dual roles, acting as a scaffold for T4P assembly and as part of an outside-in signal transduction system; the latter is a novel role for the alignment complex. Second, the observation that SadC can interact with a component of the T4P (PilO) and the flagellar motor (MotC), and both of these interactions modulate SadC activity to impact surface behaviors like swarming and biofilm formation, indicates that this DGC acts as bridge point to link surface-sensing inputs from these two motility machines. Third, our work highlights the important role of fine-tuning levels of cyclic-di-GMP and maintaining control of uniformity in signal levels for a population during early biofilm formation.

Cyclic-di-GMP heterogeneity in *P. aeruginosa* can be driven by the Wsp system or the phosphodiesterase, DipA, through the cyclic-di-GMP receptor, FimW, or through the chemotaxis machinery [18, 21, 22]. Our work here shows a variation on this theme, in that SadC’s interaction with PilO and MotC seems to restrict the temporal variation of cyclic-di-GMP levels; disrupting PilO-SadC interactions results in wider range in signal levels that impact surface behaviors.

Finally, our data indicate that the population of surface-associated cells can be partitioned into an “on” and “off” state, echoing findings of Parsek, Jenal and Miller groups [18, 21, 22]. Our data indicate that PilO-SadC interactions impact cyclic-di-GMP heterogeneity and homeostasis in two distinct but complementary ways: switching between cyclic-di-GMP “on” and “off” states as well as controlling the temporal variation in signal levels for actively signaling cells.

Our findings raise some interesting questions for future investigation. PilO interacts with both PilN [12] and SadC; what are the dynamics of these interactions? Could all three proteins form a large complex whose interaction strength varies with planktonic versus surface growth, or alternatively, could there be a surface contact-dependent partner switching mechanism? And what role does PilY1 play in the context of this model? This work grew out of the observation that surface-dependent stimulation of cyclic-di-GMP by PilY1 requires SadC [6, 23] and the alignment complex [6]. Recent exciting work by Sogaard-Andersen and colleagues showed that PilO is highly enriched in PilY1-FLAG pull-down assays and cryo-electron tomograph shows that PilY1 likely contacts the alignment complex via PilO [24], a findings consistent with our previous work that PilY1 is likely secreted through the T4P machinery [6]. How PilY1 senses surface contact, the role of the putative mechanosensitive von Willibrand A (vWA) domain of PilY1 in surface sensing, and how any such signal is transduced are all still open questions.

## Materials and Methods

Detailed Materials and Methods can be found in the Supplementary Material.

### Bacterial strains, plasmids, media and growth conditions

PA14-UCBPP [25] was used as a WT *P. aeruginosa* and *E. coli* S17 was used for chromosomal mutations and BiFC analysis throughout. Strains are listed in S1 Appendix, Table S2; plasmids are in S1 Appendix, Table S3; and primers are in S1 Appendix, Table S4. All strains were routinely grown in 5 ml lysogeny broth (LB) medium and maintained on 1.5% LB agar plates with appropriate antibiotics, as necessary [6, 10]. Biofilm, swarming and twitching assays [26–28], as well as flow cells [18–20] and bacterial adenylate cyclase two hybrid assays [29–31] were performed and quantified as reported with additional details outlined in the Supplementary Methods.

### Molecular and biochemical methods

All in frame deletions and chromosomal point mutations were generated at the native locus. Plasmid and mutant construction [32, 33], Western blot analysis and quantification [34], sub-cellular fractionation of proteins [35, 36], protein concentration, and quantification of cyclic-di-GMP [10] were performed as reported.

### Imaging

Single cell tracking and quantification was performed as reported [18, 19]. We used the pCdrA::*gfp* reporter [37] to monitor cyclic-di-GMP levels of surface-attached cells.

## Acknowledgements

We thank the Institute of Biomolecular Targeting (BioMT COBRE) at Dartmouth for providing sequencing, imaging and protein analysis. The BioMT COBRE is supported by NIH award P20-GM113132. We also thank Lijun Chen at Michigan State University for mass spectrometry analysis and Lori Burrows for supplying the PilO antibody. We would also like to thank Zdenek Svindrych for help with image analysis for BiFC experiments. This work was supported by the NIH via awards R37 AI83256 to GAO and R01AI43730 to GW. WCS is funded by a National Science Foundation Graduate Research Fellowship under DGE-1650604 and DGE-2034835.

## References

1. O’Toole, G.A. and G.C. Wong, Sensational biofilms: surface sensing in bacteria. Curr Opin Microbiol, 2016. 30: p. 139–146.

2. Ellison, C.K., et al., Obstruction of pilus retraction stimulates bacterial surface sensing. Science, 2017. 358(6362): p. 535–538.

3. Persat, A., et al., Type IV pili mechanochemically regulate virulence factors in Pseudomonas aeruginosa. Proc Natl Acad Sci U S A, 2015. 112(24): p. 7563–8.

4. Utada, A.S., et al., Vibrio cholerae use pili and flagella synergistically to effect motility switching and conditional surface attachment. Nat Commun, 2014. 5: p. 4913.

5. Conrad, J.C., et al., Flagella and pili-mediated near-surface single-cell motility mechanisms in P. aeruginosa. Biophys J, 2011. 100(7): p. 1608–16.

6. Luo, Y., et al., A hierarchical cascade of second messengers regulates Pseudomonas aeruginosa surface behaviors. mBio, 2015. 6(1).

7. Gibiansky, M.L., et al., Bacteria use type IV pili to walk upright and detach from surfaces. Science, 2010. 330(6001): p. 197.

8. Hug, I., et al., Second messenger-mediated tactile response by a bacterial rotary motor. Science, 2017. 358(6362): p. 531–534.

9. McCarter, L. and M. Silverman, Surface-induced swarmer cell differentiation of Vibrio parahaemolyticus. Mol Microbiol, 1990. 4(7): p. 1057–62.

10. Baker, A.E., et al., Flagellar stators stimulate c-di-GMP production by Pseudomonas aeruginosa. J Bacteriol, 2019. 201(18).

11. Kuchma, S.L., et al., Cyclic-di-GMP-mediated repression of swarming motility by Pseudomonas aeruginosa: the pilY1 gene and its impact on surface-associated behaviors. J Bacteriol, 2010. 192(12): p. 2950–64.

12. Leighton, T.L., et al., Novel role for PilNO in Type IV pilus retraction revealed by alignment subcomplex mutations. J Bacteriol, 2015. 197(13): p. 2229–2238.

13. Tammam, S., et al., PilMNOPQ from the Pseudomonas aeruginosa type IV pilus system form a transenvelope protein interaction network that interacts with PilA. J Bacteriol, 2013. 195(10): p. 2126–35.

14. Zhu, B., et al., Membrane association of SadC enhances its diguanylate cyclase activity to control exopolysaccharides synthesis and biofilm formation in Pseudomonas aeruginosa. Environ Microbiol, 2016. 18(10): p. 3440–3452.

15. Kelley, L.A., et al., The Phyre2 web portal for protein modeling, prediction and analysis. Nat Protoc, 2015. 10(6): p. 845–58.

16. Mudumbi, K.C., et al., The pathogenic A391E mutation in FGFR3 induces a structural change in the transmembrane domain dimer. J Membr Biol, 2013. 246(6): p. 487–93.

17. Li, E., W.C. Wimley, and K. Hristova, Transmembrane helix dimerization: beyond the search for sequence motifs. Biochim Biophys Acta, 2012. 1818(2): p. 183–93.

18. Armbruster, C.R., et al., Heterogeneity in surface sensing suggests a division of labor in Pseudomonas aeruginosa populations. Elife, 2019. 8:e59154.

19. Lee, C.K., et al., Multigenerational memory and adaptive adhesion in early bacterial biofilm communities. Proc Natl Acad Sci U S A, 2018. 115(17): p. 4471–4476.

20. Lee, C.K., et al., Social cooperativity of bacteria during reversible surface attachment in young biofilms: a quantitative comparison of Pseudomonas aeruginosa PA14 and PAO1. mBio, 2020. 11:e026440–19.

21. Kulasekara, B.R., et al., c-di-GMP heterogeneity is generated by the chemotaxis machinery to regulate flagellar motility. Elife, 2013. 2: p. e01402.

22. Laventie, B.J., et al., A surface-induced asymmetric program promotes tissue colonization by Pseudomonas aeruginosa. Cell Host Microbe, 2019. 25(1): p. 140–152.

23. Kuchma, S.L., E.F. Griffin, and G.A. O’Toole, Minor pilins of the type IV pilus system participate in the negative regulation of swarming motility. J Bacteriol, 2012. 194(19): p. 5388–403.

24. Treuner-Lange, A., et al., PilY1 and minor pilins form a complex priming the type IVa pilus in Myxococcus xanthus. Nat Commun, 2020. 11(1): p. 5054.

25. Rahme, L.G., et al., Common virulence factors for bacterial pathogenicity in plants and animals. Science, 1995. 268(5219): p. 1899–902.

26. Ha, D.G., S.L. Kuchma, and G.A. O’Toole, Plate-based assay for swarming motility in Pseudomonas aeruginosa. Methods Mol Biol, 2014. 1149: p. 67–72.

27. O’Toole, G.A., Microtiter dish biofilm formation assay. J Vis Exp, 2011 (47).

28. O’Toole, G.A. and R. Kolter, Flagellar and twitching motility are necessary for Pseudomonas aeruginosa biofilm development. Mol Microbiol, 1998. 30(2): p. 295–304.

29. Karimova, G., et al., A bacterial two-hybrid system based on a reconstituted signal transduction pathway. Proc Natl Acad Sci U S A, 1998. 95(10): p. 5752–6.

30. Jh, M., A short course in bacterial genetics. Cold Spring Harbor Press, 1992.

31. Smale, S.T., Beta-galactosidase assay. Cold Spring Harb Protoc, 2010. 2010(5): p. pdb prot5423.

32. Choi, K.H., A. Kumar, and H.P. Schweizer, A 10-min method for preparation of highly electrocompetent Pseudomonas aeruginosa cells: application for DNA fragment transfer between chromosomes and plasmid transformation. J Microbiol Methods, 2006. 64(3): p. 391–7.

33. Bachman, J., Site-directed mutagenesis. Methods Enzymol, 2013. 529: p. 241–8.

34. Bradford, M.M., A rapid and sensitive method for the quantitation of microgram quantities of protein utilizing the principle of protein-dye binding. Anal Biochem, 1976. 72: p. 248–54.

35. Nunn, D.N. and S. Lory, Cleavage, methylation, and localization of the Pseudomonas aeruginosa export proteins XcpT, -U, -V, and -W. J Bacteriol, 1993. 175(14): p. 4375–82.

36. Hinsa, S.M. and G.A. O’Toole, Biofilm formation by Pseudomonas fluorescens WCS365: a role for LapD. Microbiology (Reading), 2006. 152(Pt 5): p. 1375–83.

37. Rybtke, M.T., et al., Fluorescence-based reporter for gauging cyclic di-GMP levels in Pseudomonas aeruginosa. Appl Environ Microbiol, 2012. 78(15): p. 5060–9.

