## Supplemental Figures for "Fine tuning cyclic-di-GMP signaling in *Pseudomonas aeruginosa* using the type 4 pili alignment complex"

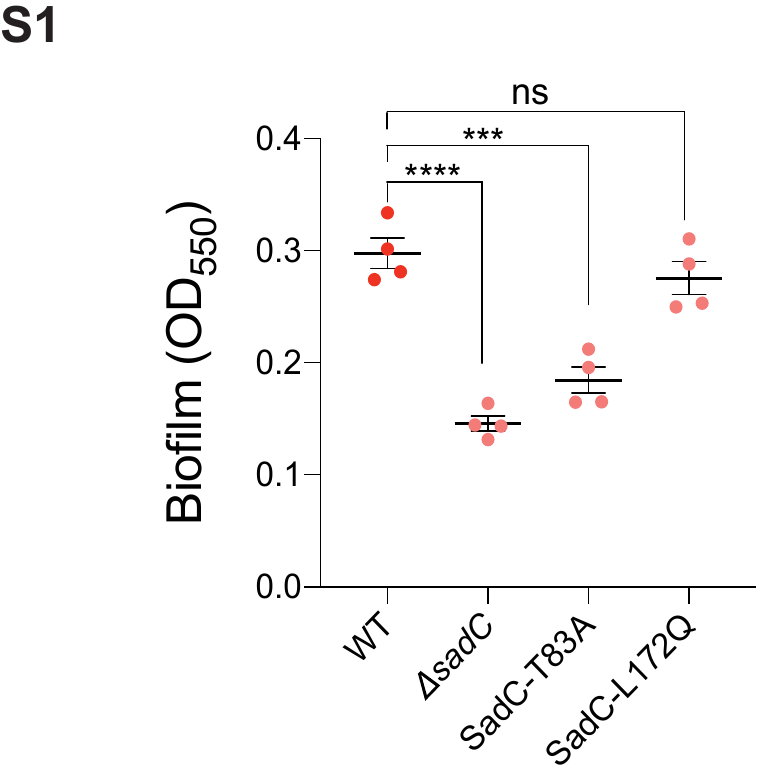

**Figure S1. Biofilm formation assays.** Absorbance measurements at OD_550_ for static crystal violet-quantified biofilm assay for WT and *sadC* variants. Biofilm formation was assessed after growth at 37 ^o^C for 24 h in biofilm medium as described in Materials and Methods. Data represents four biological replicates and statistical significance was determined using one-way ANOVA and a Dunnets post-hoc test. p-values: p ≤ *** 0.0001, p ≤ ****0.0001; ns, not significant.

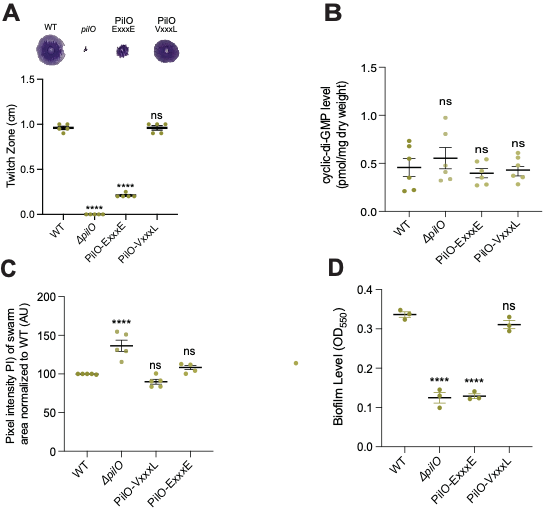

**Figure S2. Twitching motility, and quantification of cyclic-di-GMP and cyclic-di-GMP related behaviors for PilO protein mutants in *P. aeruginosa*.** **A.** Representative images (top) and quantification of twitch zones (bottom) following stabs on LB agar plates and incubation at 37 ^o^C for 24 h. **B.** Quantification of global pools of cyclic-di-GMP normalized to dry weight for WT and PilO variants. Nucleotides were extracted from cells scraped from swarm plates that were incubated 37 ^o^C for 14 h. **C.** Swarm assays. Pixel intensity (PI) of swarm area for WT, Δ*pilO*, PilO-ExxxE and PilO-VxxxL strains after incubation on 0.52% swarm medium at 37 ^o^C for 14 h. WT swarm area is normalized to 100%. **D.** Absorbance at OD_550_ for biofilm assay after incubation in swarm medium (described in Materials and Methods). Dots on each graph represent the total number of biological replicates. In all panels, error bars are SEM and p-value was determined by one-way ANOVA and a Dunnets post-hoc test. p-values: p ≤ ***** 0.0001; ns, not significant.

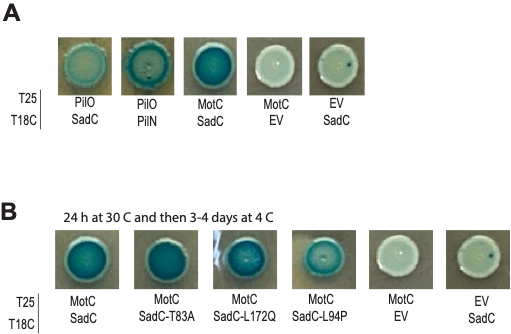

**Figure S3. BACTH analysis.** Representative spots of images from BACTH assay with co-transformations of the indicated strains fused to the T25 and T18 fragments from the adenylate cyclase from *Bordetella pertussis*. Empty vector with MotC and SadC are included as the negative controls.

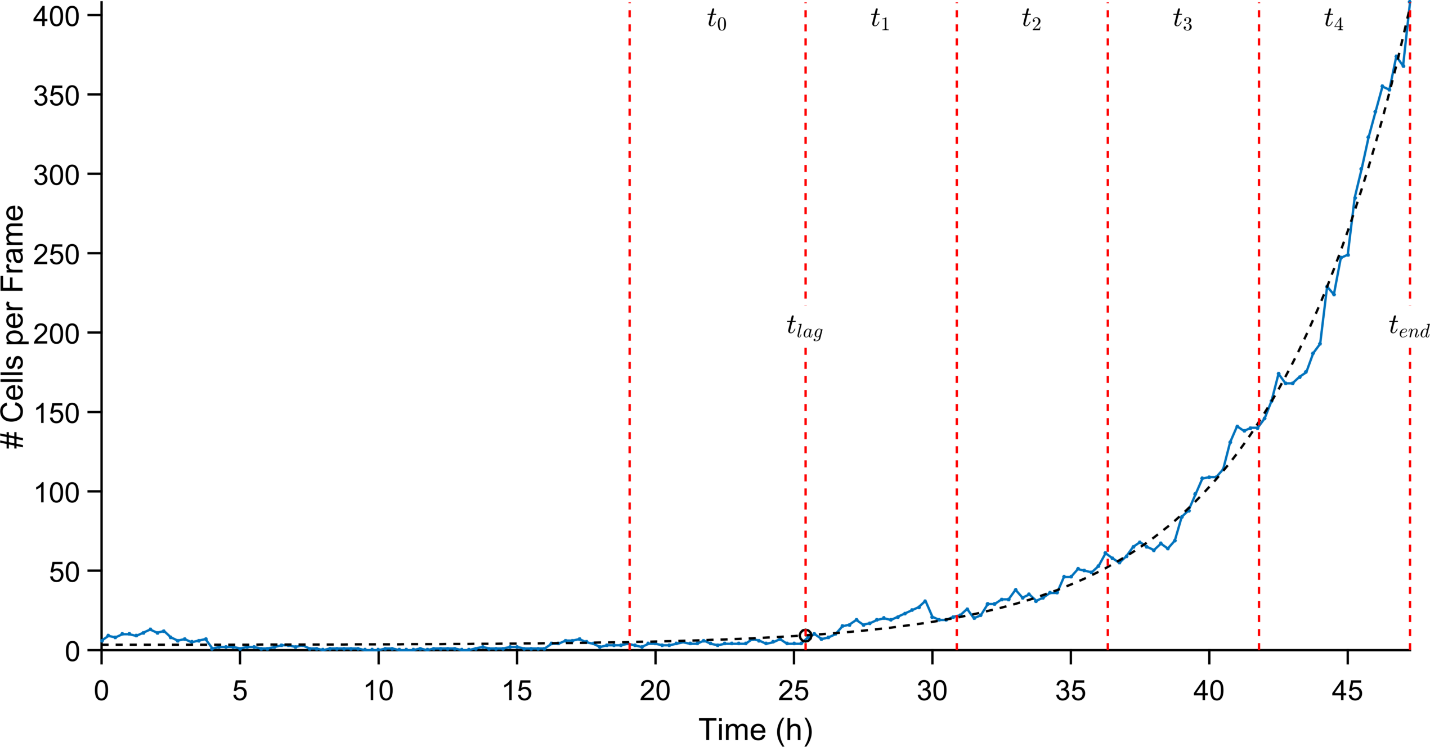

**Figure S4. Method for analyzing time course for surface population.** Representative surface exponential growth curve for bacterial cells as they transition from reversible to irreversible attachment. The growth curve is defined as the count of bacteria on the surface as a function of experiment time. Time points $t_{0}$ to $t_{4}$ are obtained from fitting the surface growth curves to the function $N_{e}\left( t \right)=N_{e}\left( 0 \right)\exp\left( \frac{t-t_{lag}}{\tau_{e}} \right)+N_{c}$where $N_{e}\left( 0 \right)$ is the number of cells at time 0, $t_{lag}$ characterizes the time scale of the lag period during which $N_{e}$ is roughly constant and after which $N_{e}$ increases exponentially, $N_{c}$ is the average number of cells present on the surface during the lag period, and $\tau_{e}$ characterizes the time scale of exponential increase. $t_{lag}$ and $t_{end}$ are then used to define $t_{0}$ to $t_{4}$ as follows:

$$t_{0}=\frac{3}{4}t_{lag}<t\leq t_{lag}$$

$$t_{1}=t_{lag}<t\leq t_{lag}+\frac{1}{4}\left( t_{end}-t_{lag} \right)$$

$$t_{2}=t_{lag}+\frac{1}{4}\left( t_{end}-t_{lag} \right)<t\leq t_{lag}+\frac{1}{2}\left( t_{end}-t_{lag} \right)$$

$$t_{3}=t_{lag}+\frac{1}{2}\left( t_{end}-t_{lag} \right)<t\leq t_{lag}+\frac{3}{4}\left( t_{end}-t_{lag} \right)$$

$$t_{4}=t_{lag}+\frac{3}{4}\left( t_{end}-t_{lag} \right)<t\leq t_{end}$$

Temporal progressions in cyclic-di-GMP for WT, *pilO* and *sadC* variants are analyzed at these five time points (t_0_- t_4_) for the data shown in Figure 4.

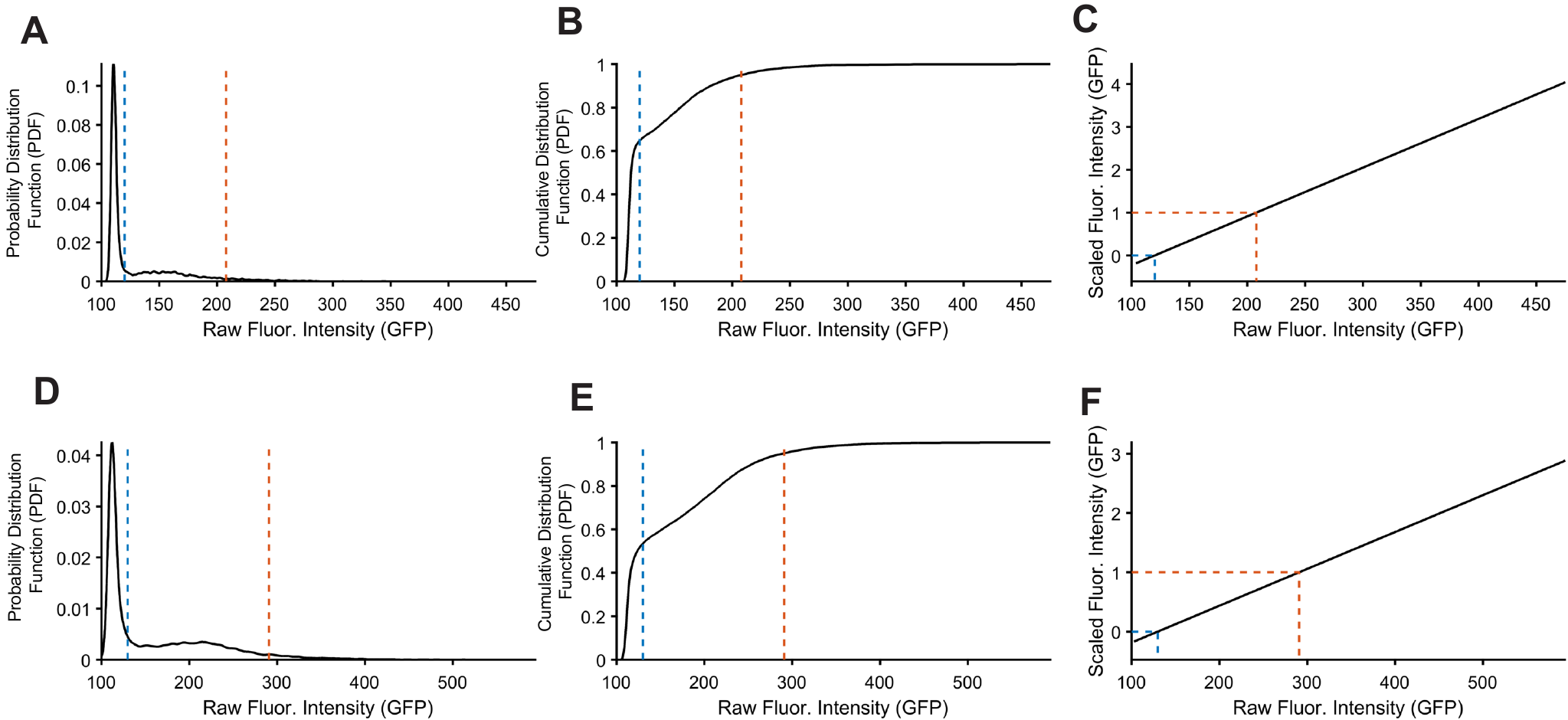

**Figure S5.** Normalizing fluorescence intensities for single cell tracking across different microscope setups (top and bottom row). For each of the two microscope setups, we look at the intensity distribution for WT. **A, B.** For the probability distribution function (PDF), there is a bimodal distribution: one sharp peak at lower intensity and one broad peak at higher intensity. A cutoff intensity value was manually picked in between these two peaks (blue dashed lines). **C, D**. This cutoff was confirmed by plotting the cumulative distribution function (CDF) and seeing that the cutoff lies on the cusp where the CDF changes slopes (a steeper slope in the CDF represents a sharper peak in the PDF, while a shallower slope represents a broader peak). To linearly rescale the raw intensities from the different setups onto the same scale, we need 2 points of reference. One is the cutoff (blue dashed lines), which we set the resulting scaled fluorescence intensity to 0. The second point is the 95^th^ percentile (red dashed lines), which we set the resulting scaled fluorescence intensity to 1. With these points of reference, we then linearly rescale the raw intensities to obtain scaled fluorescence intensities (**E, F**), which we then use for subsequent analyses and specifically for the data presented in Figure 4.
