## Supplemental Material for "Fine tuning cyclic-di-GMP signaling in *Pseudomonas aeruginosa* using the type 4 pili alignment complex"

**Novel role of the T4P alignment complex in tuning cyclic-di-GMP signaling in *Pseudomonas aeruginosa***

Running Title: T4P alignment complex and surface sensing

^*^Corresponding author:

George A. O’Toole

Dept. of Microbiology & Immunology

Rm 202 Remsen Building

66 College Street

Geisel School of Medicine at Dartmouth

Hanover, NH 03755

**This PDF file includes:**

Supplemental Materials and Methods

Table S1-S3

SI References

**Materials and Methods**

**Bacterial strains, plasmids, media and growth conditions**

All bacterial strains used in this study are listed in the supplementary material in **Table S1**. *Pseudomonas aeruginosa* PA14-UCBPP, *Escherichia coli* S17- λpir and *E. coli* BTH101 strains were routinely grown in 5 ml lysogeny broth (LB) medium and maintained on 1.5% LB agar plates with appropriate antibiotics if necessary. Overnight cultures were grown in LB with shaking at 220 rpm on a roller drum. *Saccharomyces cerevisae* InvSc1 (Thermo Fisher) used for cloning was maintained on yeast peptone dextrose (YPD - 1% Bacto yeast extract, 2% Bacto peptone, and 2% dextrose) with 2% agar. Twenty-five microgram per milliliter (25 µg/ml) of gentamycin (GoldBio) for *P. aeruginosa* and 10 µg/ml for *E. coli* was used when selection was required. Kanamycin (GoldBio) and carbenicillin (GoldBio) were used at 50 µg/ml for *E. coli.* 5-bromo-4-chloro-3-indolyl-ß-D-galactopyranoside (x-gal; GoldBio) and Isopropyl β-D-1-thiogalactopyranoside (IPTG; GoldBio) were used at 40 µg/ml and 0.5 mM concentrations respectively for *E. coli* BTH101 cells. All plasmids and oligonucleotides used in this study are listed in **Tables S2 and S3**.

**Plasmid Construction**

All chromosomal in-frame gene deletions were constructed using the pMQ30 shuttle vector carrying the flanking regions of the gene via homologous recombination [1] . *S. cerevisae* was grown overnight at 30 ^o^C in YPD. Synthetic defined medium without uracil (Sunrise Science Products) was used to select for yeast colonies with the plasmid construct. Plasmids were extracted from yeast using the ‘smash and grab’ method and transformed by electroporation into *E. coli S17* cells and grown on LB plates with 10 µg/ml gentamycin at 30 ^o^C overnight [2]. Colony polymerase chain reaction (PCR) amplification and sequencing was used to confirm plasmid construction. Plasmids were introduced in *P. aeruginosa* by conjugation and merodiploids were selected on 25 µg/ml gentamycin and 20 µg/ml nalidixic acid after which cells were counter-selected on LB with 10% sucrose-containing medium with no added salt [3]. Deletions were confirmed by colony PCR amplification and sequencing with primers flanking the gene. All sequencing was done at the Molecular Biology Core at the Geisel School of Medicine at Dartmouth.

**Construction of chromosomal point mutations**

Point mutations in the *pilO* and *sadC* genes were made using a modified *in vitro* site-directed mutagenesis protocol [4]. Forward and reverse complementary primers consisting of the nucleotide codon sequence encoding for the mutation of interest were used to separately amplify the pMQ30 (for chromosomal mutations) or pkT25/pUT18C (for BACTH studies) parental plasmids with the gene of interest using high fidelity Phusion polymerase (NEB). After four cycles of amplification, the products of these reactions were combined and amplified for an additional 18 cycles with additional Phusion polymerase added. The parental plasmid was digested for 4 h using Dpn1 (NEB) at 37 ^o^C. Following digestion, the PCR product was transformed into *E. coli S17* cells and selected on LB with appropriate antibiotics*.* Plasmid containing the desired point mutation was isolated and confirmed by sequencing. Introduction of mutations on the chromosome was done by conjugation and counter-selection as described above. All chromosomal mutations were verified by PCR amplification and sequencing.

**Construction of SadC-FLAG plasmid**

To build SadC-3xFLAG constructs, we used the 375 amino acid sequence for the full length SadC protein (GenBank entry ABJ13602) and the numbering of all mutations in this manuscript is based on this sequence. The amino acid sequence of SadC is based on the annotation of the PA14_56280 open reading frame (ORF) in the *P. aeruginosa* PA14 genome as posted on pseudomonas.com. It should be noted that the PA14_56280 ORF is predicted to translate a 375 amino acid protein, however, there is a 111 amino acid sequence upstream of this, ORF that includes a stop codon at the 86^th^ amino acid; this 111 amino acid sequence was therefore not included in the design of all FLAG tag constructs. A G-block containing a linker and 3x-FLAG tag sequence was ordered from IDT and cloned into pmq30 at the C terminus of full length SadC (**Table S1, S2**). SadC-3xFLAG was introduced at the native locus of *P. aeruginosa* by homologous recombination.

**Construction of PilN-PilO_TM_ chimera**

To construct the PilN-PilO_TM_ chimera**,** the transmembrane of *pilN* (residues 22-43) was replaced with that of PilO (residues 23-45). The transmembrane regions of each protein were determined using the open-source tool, Protter and from work published by the Burrows group [5]. PilN-PilO_TM_ chimera was fused to the C-terminus of the T25 fragment in the pKT25 plasmid for BACTH analysis. Synthesis of the chimera and cloning were done by Fisher.

**Bacterial adenylate cyclase two hybrid (BACTH) assay**

Full length genes and gene variants (*sadC, pilN* and *pilN-pilO_TM_* chimera) were cloned into the pKT25 and pUT18C expression vectors by restriction digest and ligation which generated C-terminal fusions to both the T25 and T18 fragments. These protein fusions were chosen in order to preserve the catalytic activity of SadC and to ensure proper localization of the fragments to the cytoplasm of PilO. To assess interaction between proteins, *E.coli* BTH101 cells were co-transformed by electroporation with each vector and allowed to recover in 500 µl LB for 1 h at 37 ^o^C based on a previously described protocol [6]. Transformations were spotted on LB plates supplemented with kanamycin, carbenicillin, IPTG and x-gal and incubated at 30 ^o^C for 40 h and then three additional days at 4 ^o^C to allow for further color development.

**Utilization of the BACTH assay to screen for mutations with altered interactions**

To identify variants of the PilO-AxxxA motif resulting in strong interaction with SadC, we used a primer-based approach combined with the site-directed mutagenesis procedure described above to specifically target the Ala residues at positions 40 and 44 in PilO. Forward and reverse complementary primers consisting of random base pairs, achieved by adding NNN at positions coding for the Ala residues in the AxxxA motif, were used to amplify T25-PilO parental plasmid. Following rounds of PCR and parental plasmid digestion, the final product containing the mutant library was co-transformed with the T18C-SadC vector and plated for single colonies on LB medium supplemented with the appropriate antibiotics, IPTG and x-gal. Colonies with a dark blue color were re-patched, and the *pilO*-containing plasmid isolated and sequenced to determine the residue changes. Co-transformation of the T25-PilO and T18C-SadC fragments were included as a positive control and a comparison for blue color intensity.

**β-galactosidase activity assay**

*E.coli* BTH101 cells with co-transformed plasmids were plated on the same LB medium as used for the blue/white BACTH plate read out but without x-gal. Cells were scraped from the plates following incubation after 40 h and lysed with 25 µl 0.1% SDS, 50 µl chloroform followed by 10 s vortexing and 5 min incubation at 30 ^o^C . β-galactosidase activity was calculated using the following equation:

$$Miller units= \frac{{OD}_{400}-1.75*{OD}_{550}}{time in mins*{OD}_{600}*volume of cells} x 1000$$

as described by Miller [7, 8].

**Bimolecular fluorescence complementation (BiFC)**

*E. coli* S17 strains co-transformed with pRSF-Duet plasmid with PilO and SadC YFP fusions were grown overnight in LB at 30 ^o^C. Six microliters of the overnight culture were transferred to 2.5 mL fresh LB and sub-cultured with 0.5 mM IPTG in a six well plate (Falcon) and incubated at 30 ^o^C in a floor rotary shaker (New Brunswick Scientific) with shaking at 100 rpm. For image analysis, 100 µl of cells were pelleted and washed with PBS and imaged on a wet mount.

**Image acquisition and data analysis**

All images were acquired using a Nikon Eclipse Ti inverted microscope equipped with a Hamamatsu ORCA-Flash 4.0 camera and imaged through either a Plan Apochromat 100x Ph3 Oil or Plan Fluor 40x DIC M N2 objective. All images were acquired using a Nikon Eclipse Ti inverted microscope equipped with a Hamamatsu ORCA-Flash 4.0 camera and images through either a Plan Apochromat 100x Oil or Plan Fluor 40x DIC M N2 objective. Fluorescence intensity of the individual bacterial cells was measured using a custom ImageJ macro. Specifically, the masks corresponding to the individual cells were generated by thresholding the background-corrected phase contrast images (the cells appearing dark on a bright background). These masks were then overlaid on the corresponding background-corrected epifluorescence images and the mean fluorescence of the individual cells. Images are from at least two biological on different days with at least 5 fields for each strain per biological replicate. Statistical analysis was performed with a Mann U test.

**Swarm motility assay and quantification**

Swarming motility assays were performed as described by Ha and O’Toole [9]. Swarm medium consisted of 0.52% agar plates containing M8 minimal medium (Na_2_HPO_4_, KH_2_PO_4_ NaCl) supplemented with glucose (0.2% v/v), casamino acids (0.5% v/v) and MgSO_4_ (1 mM). Swarm plates were inoculated with 2.5 µl of overnight culture and incubated upright at 37^o^C for 14 h. Pictures were taken of swarm plates and “Total Swarm Area” which is a measure of the pixel intensity (PI) was analyzed using ImageJ by first selecting the swarm area, converting images to grayscale  (Image → Type → 8-bit), thresholding the image and then analyzing the number of pixels in the swarm using the default settings.

**Biofilm assay**

Overnight cultures (1.5 µl) were inoculated in U-bottom 96 well plates (Costar) containing 100 µl swarming medium and incubated at 37^o^C for 24 h. Plates were then stained with 100 µl 0.1% crystal violet for 20 mins at room temperature and de-stained for 20 mins with 125 µl de-staining solution (40% glacial acetic, 40% methanol and 20% H_2_O v/v). Absorbance was read at OD_550_ and destaining solution was included as the blank. Biofilm assays were done similar to published work by the O’Toole group [10, 11].

***In vivo* cyclic-di-GMP quantification**

Nucleotides were extracted from *P. aeruginosa* cells scraped from swarm plates after incubation for 37^o^C for 14 h. Cells were removed from plates by gently scraping with a cover slip to avoid scraping the agar and immediately placed on ice. Cell pellets were re-suspended in 250 µl nucleotide extraction buffer (methanol/acetonitrile/dH_2_O 40:40:20 + 0.1 N formic acid) and incubated at -20^o^C for 1 h. Following nucleotide extraction, cells were spun for 5 mins at 4^o^C and 200 µl of supernatant was removed and added to 8 µl of 15% NH_4_HCO_3_ stop solution. Nucleotides were dried in a speed vacuum and resuspended in 200 µl HPLC grade water (JT Baker) and placed in screw cap vials (Agilent Technologies). Quantification of cyclicyclic-di-GMP levels was done by liquid chromatography-mass spectrometry (LC-MS/MS) at the Mass Spectrometry Facility at Michigan State University. All samples were normalized to dry weight and expressed as $\frac{pmol}{mg dry weight}$.

**Sub-cellular protein fractionation**

Overnight cultures of WT, Δ*pilO*, PilO-ExxxE and PilO-VxxxL were grown in LB at 37 ^o^C. Cells were diluted in 1:10 in 100 mL swarm medium (as described above) and sub-cultured at 37 ^o^C with shaking at 250 rpm for 3-4 h until OD_600_ is ~0.6. Cells were harvested by centrifuging at 5000 rpm at 4 ^o^C for 15 mins. Supernatant was removed and pellets were resuspended in 3 mL Buffer C (50mM Tris-HCl pH 8.0, 1 mM EDTA, 2 mM MgCl_2_, freshly made 1X protease inhibitors EDTA-free tablets – Sigma) with 1 μl Benzonase (Novagen). Cells were by lysed using a French Press and a sonicator (Fischer Scientific Sonic Dismembrator Model 500) with the following settings: pulse time 10 s, amplitude-30, for one cycle. Following lysis, samples were incubated on ice for 5 mins and then spun at low speed at 13 000 rpm for 4 ^o^C for 5 mins to pellet unbroken cells. An aliquot of the supernatant designated as the whole cell was removed and stored at -20 ^o^C until further analysis. The remaining supernatant was transferred to microfuge tubes (Beckman Coulter) and spun at 50, 000 x g at 4^o^C for 1 h in the ultracentrifuge (Beckman TL-100) to separate the cytoplasmic fraction (supernatant) from the total membrane fraction (pellet). Most but not all of the supernatant was collected to avoid contamination with the pellet. Supernatants were stored at -20 ^o^C. The pellet was washed in 0.5 mL Buffer B (20mM Tris-HCl, pH7.6, 1X fresh protease inhibitor) by gently inverting the tubes several times followed by a brief spin. The supernatant was removed and pellets were resuspended in 0.1 mL Buffer B with glycerol (10% final concentration) and sarkosyl (2% final concentration) and vigorously vortexed for three rounds of 30 s vortex and 30 s on ice. To solubilize membranes, samples were incubated overnight on ice in the cold room and the following day at room temperature for 20 mins on an inverting rotator. Solubilized membranes were transferred to new microfuge tubes and spun 100,000 x g at 4 ºC for 1 h. The inner-membrane fraction was collected as the supernatant and stored until further analysis. Twenty five micrograms of total protein were ran on a 12% Tris-HCl Precast SDS-PAGE gel (Bio-Rad) for whole cell lysate, inner-membrane and cytoplasmic fractions during Western blot analysis. SecY (~50 kD) and Cas3 (~120 kD) which are known to localize to the inner-membrane and cytoplasm, respectively, were blotted for and used as controls for each fraction described in other studies from our group [12, 13].

**Protein quantification**

Total protein levels in whole cell lysate, inner-membrane and cytoplasm was quantified using the Bio-Rad protein assay Dye Reagent as per the manufacturer’s instructions as outlined by Bradford [14].

**Western blot analysis for PilO protein levels and SadC-FLAG**

Chromosomal FLAG-tagged SadC of WT, SadC-T83A, SadC-L172Q and *ΔpilO* were grown overnight in LB at 37 ^o^C. Cells were normalized to OD600 = 2 and lysed with sample buffer with β-mercaptoethanol. Samples were ran on a 10% Tris-HCl Precast SDS-PAGE gel (Bio-Rad) and blotted onto 0.45 µm pore size nitrocellulose membrane (Bio-Rad) using the 1.5 mm pre-programmed method on a Trans-Blot Turbo Transfer System (Bio-Rad). The membrane was incubated in blocking buffer (LI-COR Blocking Buffer in TBS) for 1 h at room temperature, washed in TBST_0.1%_ for 5 mins x3 and incubated for 1 h in monoclonal anti-FLAG M2 antibody (Sigma) (1:10,000 dilution) in TBST_0.1%_ buffer. Following incubation with primary antibody, the membrane was washed in TBST_0.1%_ for 5 mins x3 and incubated for 1 h with goat anti-mouse in TBST_0.1%_ (1:15,0000 dilution) secondary antibody. Incubation with secondary antibody and all subsequent steps were performed in the dark. After incubation with the secondary antibody, the membrane was washed in TBST_0.1%_ for 5 mins x2 and then once in TBS. The membrane was imaged using the LI-COR Odyssey CLx imager at BioMT Core at the Geisel School of Medicine at Dartmouth. Protein levels were quantified relative to the cross-reacting band at ~32 kD using the LI-COR Image Studio Lite software by drawing a rectangle of the same size around each band and using the following background settings: average, border width of 3, segment = all.

**Strains and growth conditions for flow cell experiments**

*Pseudomonas aeruginosa* PA14 WT and its isogenic strains with mutation in the *pilO* gene (Δ*pilO*, PilO VxxxL, and PilO ExxxE mutations) were used in this study. For cyclic di-GMP measurements, a plasmid-based, cyclic-di-GMP responsive transcriptional reporter, pCdrA::*gfp* [15]. Overnight cultures were diluted 1:50 and sub-cultured in swarm medium to OD_600_ of ∼0.4. Cultures were further diluted to OD_600_ of ∼0.01 to 0.03 in flow-cell medium, which consisted of M63 supplemented with 1 mM MgSO_4_, 0.05% glucose, and 0.125% CAA. The diluted cultures were then used for injection into the flow chamber.

**Flow cells**

Flow cells were purchased from Ibidi (sticky-Slide *VI*^0.4^ with a glass coverslip) and prepared as previously described [16-18]. The prepared flow cell was connected to a syringe through a 0.22-μm filter (Fisher Scientific) using Silastic silicon tubing of inner diameter 1.57 mm and outer diameter 3.18 mm (Dow Corning) and a natural Kynar PVDF female Luer to 1.6-mm barb adapter (Value Plastics DBA Nordson Medical). An in-line injection port (Ibidi) was used at the inlet for inoculating bacteria into the flow cell. Elbow connectors (Ibidi) were used to connect the chamber with tubing. The assembled system was flushed with 3% H_2_O_2_ at a volumetric flow rate of 25 mL/h using a syringe pump (KD Scientific or Harvard Apparatus) and allowed to sit for a total of 4 h including flushing time. The sterilized system was then flushed with autoclaved, deionized water at a flow rate of 5 mL/h using a syringe pump and allowed to sit overnight. Before inoculation of the bacteria into the flow cell, the flow-cell system was flushed with flow-cell medium at 30 mL/h. The diluted bacterial culture was injected into the flow cell and allowed to incubate for 10 to 20 min without flow on the heating stage at 30 °C. Flow was then started at 3 ml/h for the total flow time of 60~80 h.

**Data acquisition for flow cell analysis**

Images were taken using either an Andor iXon electron-multiplying charge-coupled-device (EMCCD) camera with Andor IQ software on an Olympus IX81 microscope equipped with a Zero Drift Correction autofocus system or an Andor Neo scientific complementary metal-oxide-semiconductor (sCMOS) camera with Andor IQ software on an Olympus IX83 microscope equipped with a Zero Drift Correction 2 continuous autofocus system. Both systems used a 100× objective, but the IX81 system used an additional 2× lens. Bright-field images were taken every 3 s (30-ms exposure time). For cyclic-di-GMP measurements, fluorescence images were also taken every 15 min (500 ms and 150 ms exposure times for the IX81 and IX83 systems, respectively) using a Lambda LS (Sutter Instrument) xenon arc lamp and a green fluorescent protein (GFP) filter. The total acquisition time was ∼80 h, resulting in ∼96,000 bright-field images and 320 fluorescence images for each fluorophore. The image size was 67 μm by 67 μm (1,024 by 1,024 pixels) and 133 μm by 133 μm (2,048 by 2,048 pixels) for the IX81 and IX83 systems, respectively.

**Image analysis and growth curves**

The image analysis algorithms and software are adapted from methods previously described [19-24] and written in MATLAB R2015a (MathWorks). All raw bright-field images were processed using a sequence of background correction, intensity normalization, Gaussian and edge filters, and Otsu thresholding to generate binary images, which contained main features of bacteria. The number of bacteria in each bright-field image over the time of acquisition was then plotted to obtain the surface growth curves for each of the strains. This surface growth curve was then fit to the following exponential function for number of cells present on the surface over time:

$N_{e}\left( t \right)=N_{e}\left( 0 \right)\exp\left[ \left( t-t_{\mathrm{lag}} \right)/{\tau_{e}} \right]+N_{c}$.

$N_{e}\left( 0 \right)$ is the initial number of cells on the surface at time t = 0. $t_{\mathrm{lag}}$ represents the lag period of the system, during which the number of cells on the surface ($N_{e}$) is roughly constant, and after which $N_{e}$begins to increase exponentially. $N_{c}$ is the average number of cells present on the surface through this lag period, and $\tau_{e}$is the time scale of exponential increase (ie. 1/$\tau_{e}$ is the rate of exponential increase). The various $t_{\mathrm{lag}}$ values were then normalized by the WT value.

**Table S1. The number of cells used to generate the distribution for each time point and strain is summarized as follows.**

| **Strain** | $\boldsymbol{t}_{\boldsymbol{0}}$ | $\boldsymbol{t}_{\boldsymbol{1}}$ | $\boldsymbol{t}_{\boldsymbol{2}}$ | $\boldsymbol{t}_{\boldsymbol{3}}$ | $\boldsymbol{t}_{\boldsymbol{4}}$ |
| --- | --- | --- | --- | --- | --- |
| WT | 483 | 1065 | 2737 | 7074 | 21599 |
| PilO-VxxxL | 598 | 1019 | 3382 | 13907 | 56880 |
| PilO-ExxxE | 1481 | 652 | 1523 | 5891 | 21062 |
| Δ*pilO* | 2295 | 720 | 934 | 1468 | 2313 |
| SadC-L172Q | 452 | 1679 | 4487 | 15931 | 69162 |
| SadC-T83A | 825 | 1447 | 3322 | 8588 | 24014 |

**Table S2. Strains used in this study**

| **SMC #s/Strain Name** | **Genotype/Description** | **Source** |
| --- | --- | --- |
| ***E. coli* strains** |  |  |
| S17-1 λ pir | *thi pro hsdR- hsdM+ ∆recA RP4-2::TcMu-Km::Tn* |  |
| BTH101 | F^-^, cya^-99^, *araD139*, *galE15*, *galK16*, rpsL1 (Str^r^), *hsdR2*, *mcrA1*, *mcrB1* | Euromedex |
| ***P. aeruginosa* strains** |  |  |
| 232 | PA14-UCBPP wildtype (WT) | [25] |
| 4465 | Δ*sadC* | [26] |
| 6763 | Δ*pilO* | [13] |
| 8238 | SadC-3xFLAG | This study |
| 8937 | SadC-3xFLAG T83A | This study |
| 8939 | SadC-3xFLAG L172Q | This study |
| 8941 | PilO A40E A44E | This study |
| 8943 | PilO A40V A44L | This study |
| 6763 | Δ*pilO* | [13] |
| 8929 | WT mKate pPcdrA::GFP | This study |
| 8930 | *ΔpilO* mKate pPcdrA::GFP | This study |
| 8931 | PilO-ExxxE mKate pPcdrA::GFP | This study |
| 8932 | PilO-VxxxL mKate pPcdrA::GFP | This study |
| 8933 | SadC-L172Q pPcdrA::GFP | This study |
| 8934 | SadC-T83A pPcdrA::GFP | This study |
| 8935 | *ΔsadC* pPcdrA::GFP | This study |

**Table S3. Plasmids used in this study**

| **Plasmid** | **Description** | **Source/SMC #** |
| --- | --- | --- |
| pmq30 | Shuttle vector for yeast cloning and Gram-negative allelic replacement, Gm^r^ | [1] |
|  | SadC-3xFLAG KI construct, Gm^r^ | This study/8237 |
|  | SadC-3xFLAG T83A KI construct, Gm^r^ | This study/8936 |
|  | SadC-3xFLAG L172Q KI construct, Gm^r^ | This study/8938 |
|  | PilO A40E A44E KI construct, Gm^r^ | This study/8940 |
|  | PilO A40V A44L KI construct, Gm^r^ | This study/8942 |
| pRSF Duet-1 | Protein expression plasmid |  |
| pPcdrA::GFP | Cyclic-GMP reporter | [15]/6911 |
| pKT25 | BACTH vector allowing fusion to the C-terminus of the cya T25 fragment, Kan^r^ | Euromedex |
| pUT18C | BACTH vector allowing fusion to the C-terminus of the cya T25 fragment, Kan^r^ | Euromedex |
| pKT25-PilO | Full length *pilO* cloned into pKT25, Kan^r^ | This study/7194 |
| pKT25-PilN | Full length *pilN*, Kan^r^ | This study/7192 |
| pKT25-PilN-PilO_TM_ | Full length *pilN* with its transmembrane replaced with that of *pilO* (PilN-O_TM_), Kan^r^ | This study/8944 |
| pUT18C-SadC | Full length *sadC* cloned into pUT18C, Cb^r^ | [27] |
| pUT18C-SadC (TM) | Transmembrane domain of sadC (amino acids 1-187) cloned into pUT18C, Cb^r^ | [27] |
| pUT18C-SadC (cyto) | Cytoplasmic domain of sadC (amino acids 188-375) cloned into pUT18C, Cb^r^ | [27] |
| pUT18C-SadC T83A | SadC-T83A cloned into pUT18C, Cb^r^ | This study/8945 |
| pUT18C-SadC L172Q | SadC-L172Q cloned into pUT18C, Cb^r^ | This study/8946 |
| pUT18C-PilO A40E A44E | PilO-A40E A44E cloned into pUT18C, Cb^r^ | This study/8947 |
| pUT18C-PilO A40V A44L | PilO-A40V A44L cloned into pUT18C, Cb^r^ | This study/8948 |
| pRSF-Duet-1  YC-PilO YN-SadC | YC-PilO and YN-SadC - N termini of PilO and SadC fused to YC and YN domains of YFP | This study/8949 |
| pRSF-Duet-1  YC-PilO YN-SadC-T83A | N terminus of PilO and SadC-T83A fused to YC and YN domains of YFP | This study/8850 |

**Table S4. Oligonucleotides used in this study**

| **Primer Name** | **Primer Sequence (5^’^ – 3^’^)** |
| --- | --- |
| QC PilO A40E A44E F | CGTCTGCGTGCTGCTGACCGAGGCGGTCCTGGAGCTGGG CTACAACTTCCATCTC |
| QC PilO A40E A44E R | GAGATGGAAGTTGTAGCCCAGCTCCAGGACCGCCTC GGTCAGCAGCACGCA GAC G |
| QC PilO-A40V A44L F | CATCGTCTGCGTGCTGCTGACCGTAGCGGTCCTGCTGCTGGGCTACAACTTCCATCTC |
| QC PilO-A40V A44L R | GAGATGGAAGTTGTAGCCCAGCAGCAGGACCGCTACGGTCAGCAGCACGCAGACGATG |
| QC SadC-T83A F | CCTGCGTTACGCCGATCCCAGCCTGGCCGAGCCGCAGGTGCTGGTGGCGATCGCC |
| QC SadC-T83A R | GGCGATCGCCACCAGCACCTGCGGCTCGGCCAGGCTGGGATCGGCGTAACGCAGG |
| QC SadC-L172Q F | GTGCTCGTGATGTTCATCGTAATGGTCTGGCAGAGCCTGTTCGCCAGCTACATACAGGC |
| QC SadC-L172Q R | CGCCTGTATGTAGCTGGCGAACAGGCTCTGCCAGACCATTACGATGAACATCACGAGCAC |

**References**

1.	Shanks, R.M., et al., *Saccharomyces cerevisiae-based molecular tool kit for manipulation of genes from gram-negative bacteria.* Appl Environ Microbiol, 2006. **72**(7): p. 5027-36.

2.	Hoffman, C.S. and F. Winston, *A ten-minute DNA preparation from yeast efficiently releases autonomous plasmids for transformation of Escherichia coli.* Gene, 1987. **57**(2-3): p. 267-72.

3.	Choi, K.H., A. Kumar, and H.P. Schweizer, *A 10-min method for preparation of highly electrocompetent Pseudomonas aeruginosa cells: application for DNA fragment transfer between chromosomes and plasmid transformation.* J Microbiol Methods, 2006. **64**(3): p. 391-7.

4.	Bachman, J., *Site-directed mutagenesis.* Methods Enzymol, 2013. **529**: p. 241-8.

5.	Leighton, T.L., et al., *Novel role for PilNO in Type IV pilus retraction revealed by alignment subcomplex mutations.* J Bacteriol, 2015. **197**(13): p. 2229-2238.

6.	Karimova, G., et al., *A bacterial two-hybrid system based on a reconstituted signal transduction pathway.* Proc Natl Acad Sci U S A, 1998. **95**(10): p. 5752-6.

7.	Smale, S.T., *Beta-galactosidase assay.* Cold Spring Harb Protoc, 2010. **2010**(5): p. pdb prot5423.

8.	JH, M., *A short course in bacterial genetics.* Cold Spring Harbor Press, 1992.

9.	Ha, D.G., S.L. Kuchma, and G.A. O'Toole, *Plate-based assay for swarming motility in Pseudomonas aeruginosa.* Methods Mol Biol, 2014. **1149**: p. 67-72.

10.	O'Toole, G.A. and R. Kolter, *Flagellar and twitching motility are necessary for Pseudomonas aeruginosa biofilm development.* Mol Microbiol, 1998. **30**(2): p. 295-304.

11.	O'Toole, G.A., *Microtiter dish biofilm formation assay.* J Vis Exp, 2011(47):p. 2437.

12.	Kuchma, S.L., et al., *BifA, a cyclic-Di-GMP phosphodiesterase, inversely regulates biofilm formation and swarming motility by Pseudomonas aeruginosa PA14.* J Bacteriol, 2007. **189**(22): p. 8165-78.

13.	Luo, Y., et al., *A hierarchical cascade of second messengers regulates Pseudomonas aeruginosa surface behaviors.* mBio, 2015. **6**:e02456-14.

14.	Bradford, M.M., *A rapid and sensitive method for the quantitation of microgram quantities of protein utilizing the principle of protein-dye binding.* Anal Biochem, 1976. **72**: p. 248-54.

15.	Rybtke, M.T., et al., *Fluorescence-based reporter for gauging cyclic di-GMP levels in Pseudomonas aeruginosa.* Appl Environ Microbiol, 2012. **78**(15): p. 5060-9.

16.	Lee, C.K., et al., *Multigenerational memory and adaptive adhesion in early bacterial biofilm communities.* Proc Natl Acad Sci U S A, 2018. **115**(17): p. 4471-4476.

17.	Lee, C.K., et al., *Social cooperativity of bacteria during reversible surface attachment in young biofilms: a quantitative comparison of Pseudomonas aeruginosa PA14 and PAO1.* mBio, 2020. **11**:e02644-19.

18.	Armbruster, C.R., et al., *Heterogeneity in surface sensing suggests a division of labor in Pseudomonas aeruginosa populations.* Elife, 2019. **8**:e45084.

19.	Zhao, K., et al., *Psl trails guide exploration and microcolony formation in Pseudomonas aeruginosa biofilms.* Nature, 2013. **497**(7449): p. 388-391.

20.	Gibiansky, M.L., et al., *Bacteria use type IV pili to walk upright and detach from surfaces.* Science, 2010. **330**(6001): p. 197.

21.	Conrad, J.C., et al., *Flagella and pili-mediated near-surface single-cell motility mechanisms in P. aeruginosa.* Biophys J, 2011. **100**(7): p. 1608-16.

22.	Jin, F., et al., *Bacteria use type-IV pili to slingshot on surfaces.* Proc Natl Acad Sci U S A, 2011. **108**(31): p. 12617-22.

23.	Utada, A.S., et al., *Vibrio cholerae use pili and flagella synergistically to effect motility switching and conditional surface attachment.* Nat Commun, 2014. **5**: p. 4913.

24.	Lee, C.K., et al., *Evolution of cell size homeostasis and growth rate diversity during initial surface colonization of Shewanella oneidensis.* ACS Nano, 2016. **10**(10): p. 9183-9192.

25.	Rahme, L.G., et al., *Common virulence factors for bacterial pathogenicity in plants and animals.* Science, 1995. **268**(5219): p. 1899-902.

26.	Bohn, Y.S., et al., *Multiple roles of Pseudomonas aeruginosa TBCF10839 PilY1 in motility, transport and infection.* Mol Microbiol, 2009. **71**(3): p. 730-47.

27.	Baker, A.E., et al., *Flagellar stators stimulate c-di-GMP production by Pseudomonas aeruginosa.* J Bacteriol, 2019. **201**(18).
